# Positively selected variants in functionally important regions of TLR7 in *Alouatta guariba clamitans* with yellow fever virus exposure in Northern Argentina

**DOI:** 10.1101/725333

**Authors:** Nicole S. Torosin, Timothy H. Webster, Hernan Argibay, Hebe Ferreyra, Marcela Uhart, Ilaria Agostini, Leslie A. Knapp

## Abstract

In 2007-2009, a major yellow fever virus (YFV) outbreak in Northern Argentina decimated the local howler monkey (*Alouatta*) population. We explored the relationship between Toll-like receptor (TLR) 7 and TLR8 gene variation and YFV susceptibility using samples from *Alouatta* individuals alive before the YFV outbreak, individuals that died during the outbreak, and individuals that survived the outbreak and are alive today. We measured genetic divergence between *Alouatta* YFV exposure groups and evaluated *Alouatta*-specific substitutions for functional consequences. We did not find different allele frequencies in the post-YFV exposure *Alouatta* group compared to the pre-exposure group. However, we identified three nonsynonymous variants in TLR7 in *A. guariba clamitans*. Two of these substitutions are under positive selection in functionally important regions of the gene. These unique coding differences in *A. guariba clamitans* may affect YFV resistance, but more work is necessary to fully explore this hypothesis.

## 3 Introduction

Yellow fever virus (YFV) is a single-stranded (ss) RNA virus endemic to Africa and South America that causes hemorrhagic disease (Beasley, McAuley, & Bente, 2014). It was introduced to the Americas on slave trade ships from Africa about 400 years ago (Bryant, Holmes, & Barrett, 2007). Soon after, the virus established a sylvatic cycle, meaning transmission involving non-human primates and mosquitoes (Hanley et al., 2013). Today, the virus continues to circulate among humans and non-human primates in Northern Argentina (Holzmann et al., 2010; Goenaga et al., 2012) (**Figure 1**).

**Figure 1:**
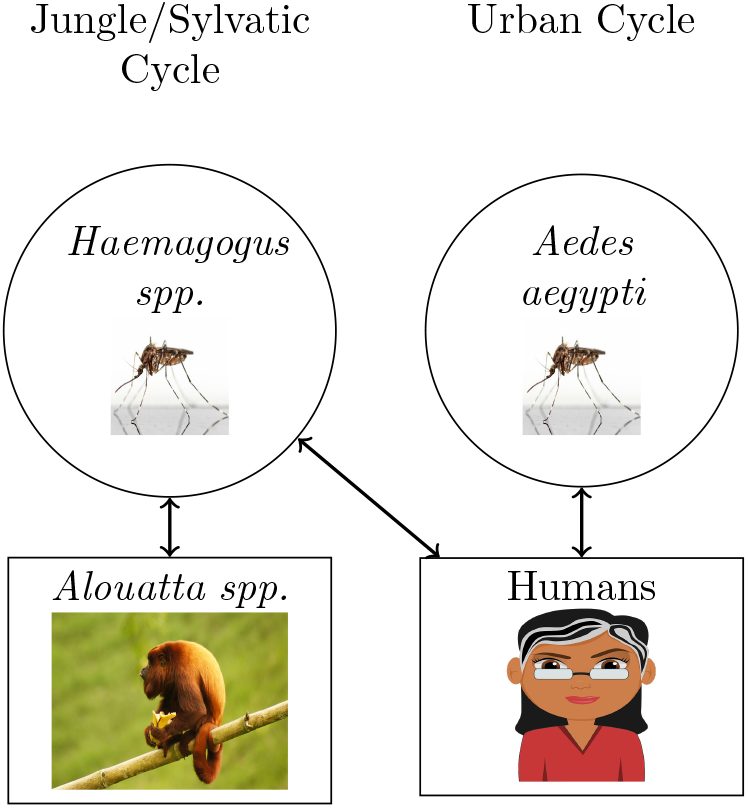
The transmission cycle of YFV between humans and howler monkeys. YFV circulates among humans through the *Aedes aegypti* mosquito vectors, called the urban transmission cycle. YFV can also be transmitted between non-human primates hosts by the *Haemagogus spp*. mosquito, this is called the sylvatic transmission cycle (Barrett & Higgs, 2007).

New World primates are generally more susceptible to YFV than Old World primates due to relatively recent exposure (Hanley et al., 2013). New World primates have likely not experienced host-pathogen interaction with YFV for enough time to evolve resistance (Hanley et al., 2013). The members of the genus *Alouatta* (howler monkeys) are particularly susceptible to this flavivirus and usually die within a week of infection (Bicca-marques & Freitas, 2010; Ministério da Saúde, 2014).

When human and non-human primate YFV transmission cycles come into contact, the risk of a sylvatic outbreak and howler monkey mortality from YFV increase. In 1966, a sylvatic YFV outbreak occurred in Argentina (Bejarano, 1979). Researchers found three dead howler monkeys and reported several human cases (Crespo, 1974; Holzmann et al., 2010). After another outbreak in 2001 in Brazil near the Argentine border, researchers found eighty deceased brown howler monkeys (Holzmann et al., 2010; Sallis, de Barros, Garmatz, Fighera, & Graça, 2003).

In 2007-2009, *Alouatta caraya* (black and gold howler) and *A. guariba clamitans* (brown howler) in Northern Argentina and Brazil suffered mass population reductions due to a YFV outbreak (Moreno et al., 2013; Holzmann et al., 2010; Bicca-marques & Freitas, 2010). This outbreak was potentially exacerbated by the deforestation of the Atlantic forest over the last 50 years (Di Bitetti, Placci, Brown, & Rode, 1994; Holzmann et al., 2010). Deforestation resulted in the intrusion of humans into the howler monkey habitat, increasing the likelihood of a YFV infected human transmitting the virus to howler monkeys (**Figure 1**) (Hanley et al., 2013). Additional factors such as non-human primate distribution, annual rainfall, and temperature can affect the chance of an outbreak as well (de Almeida et al., 2019).

The 2007-2009 outbreak provided a unique opportunity to study genetic differences between *Alouatta* individuals exposed and not exposed to YFV. We generated *A. guariba clamitans* and *A. caraya* immune gene sequences to compare genetic variants between individuals alive before the YFV outbreak, those that died during the outbreak, and individuals that survived the outbreak. We hypothesized that the surviving howler monkeys may possess advantageous genetic variants inherited from monkeys alive prior to the most recent YFV outbreak and that those that died of YFV in 2007-2009 may have lacked those variants. Alleles at greater frequency in *Alouatta* individuals alive after the 2007-2009 YFV outbreak may have been advantageous in the individual survival for this extremely susceptible species.

Previous work found positively selected genetic variants within Toll-like receptor (TLR) 7 and TLR8 in *A. guariba clamitans* and *A. caraya* that are unique to one or both species compared to Old and other New World primates (Torosin, Argibay, Webster, Corneli, & Knapp, 2019). Polymorphisms in TLR7 and TLR8 have been implicated in progression of other diseases, making these two genes strong candidates to study genetic variation that affects susceptibility to YFV (Mandl et al., 2011; Kawai & Akira, 2010; Cook, Pisetsky, & Schwartz, 2004).

## 4 Methods

### 4.1 Sample collection

This study focuses on El Parque Provincial El Piñalito in Misiones, the northernmost province of Argentina. In this park, *A. guariba clamitans* (brown howler) and *A. caraya* (black and gold howler) live sympatrically (Agostini, Holzmann, & Di Bitetti, 2010). After the YFV outbreak in 2007-2009, Holzmann and colleagues (Holzmann et al., 2010) found a total of 39 *A. caraya* and 20 *A. guariba clamitans* deceased in Misiones province. In El Piñalito, they found seven of each *Alouatta* species dead. RT-PCR for *Flavivirus* confirmed YFV infection in dead howlers in the area (Holzmann et al., 2010).

### 4.2 Sample collection

In 2017 we collected samples from taxidermied museum skins from *Alouatta* living in Misiones and the neighboring province, Corrientes, between 1949-1967. These individuals may have been exposed to YFV in the 1954 (Vezzani & Carbajo, 2008) or the 1966 outbreak (Bejarano, 1979), but since these outbreaks affected few monkeys and deforestation was not yet leading to mass exposure of human populations to *Alouatta*, we consider this possibility unlikely. To collect samples, we removed 1-1.5 cm of skin from three *A. caraya* (AC) individuals and one *A. guariba clamitans* (AGC). We stored each sample in a sterile 1.5 ml Eppendorf tube at ambient temperature until DNA extraction.

We collected liver samples from YFV positive and negative *A. caraya* deceased in Misiones between 2007-2009 as part of a YFV epizootic research project conducted by WCS-Global Health Program (**Figure 2**). We stored AC_tissue_l and AC_tissues_3-6 at −70*°C* in 100% ethanol and (AC_tissue_2) in 100% ethanol. To confirm the cause of death, we tested the samples for YFV using RT-PCR and histopathological diagnosis.

**Figure 2:**
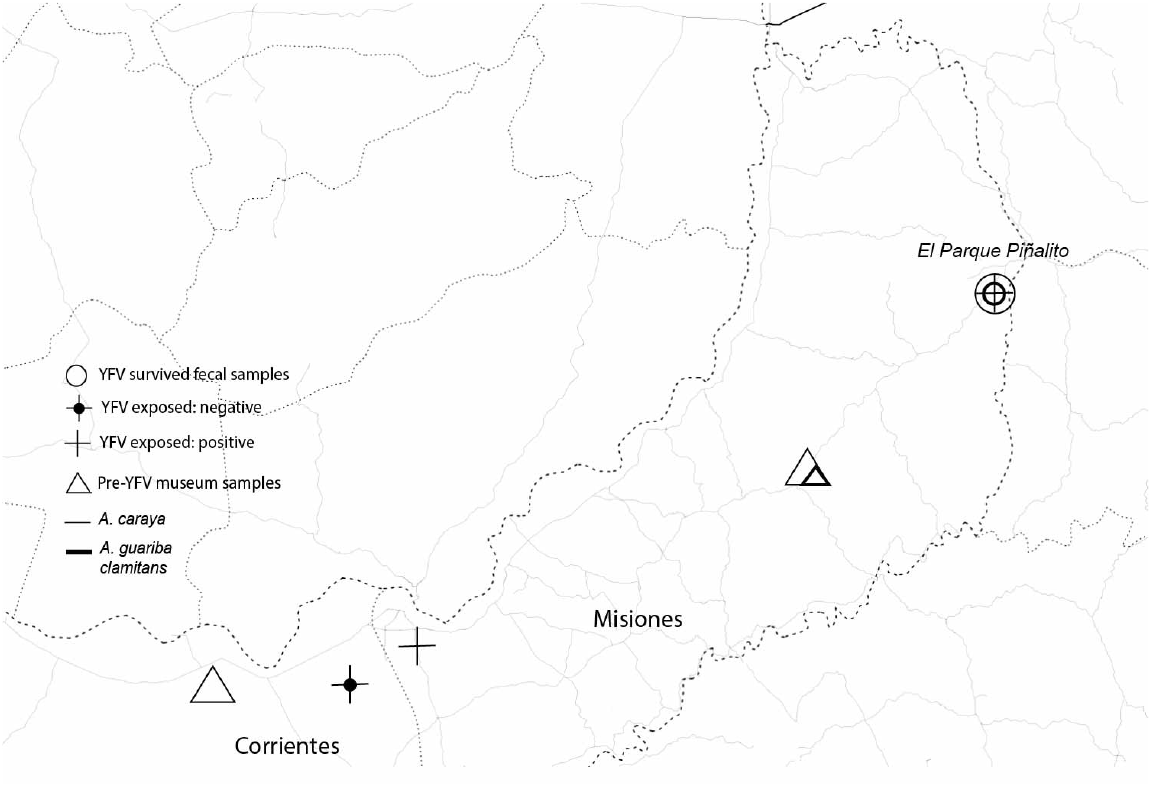
Map of samples used in this study.

We returned to El Piñalito Provincial Park, Misiones, Argentina in 2017 to determine whether any *Alouatta* were living in this region and to collect fecal samples. We collected fecal samples under contract with the Ministerio de Ecología y Recursos Naturales Renovables de Misiones Provincia, Provision number 028, file number 9910-00054/17. We placed recovered fecal pieces in sterile 15 ml collection tubes with RNAlater. We extracted DNA from fecal samples in Argentina. We used a Zymo extraction kit (Irvine, CA) with a modified protocol to target host DNA from fecal samples. We transported feces and extracted DNA to Buenos Aires under transport permit number 001494 issued by the Presidencia de la Nación Secretaría de Ambiente y Desarrollo Sustentable. We exported feces and extracted DNA from Argentina under CITES export permit 042117 and CDC import permit number 2017-03-014. We also extracted DNA from feces at the Molecular Genetics Laboratory at the University of Utah using the same Zymo extraction kit (Irvine, CA) with a modified protocol. Locations of all collected samples can be found in **Table 1**.

**Table 1:** GPS location for each sample

| **Sample** | **Latitude** | **Longitude** | **Notes** |
| --- | --- | --- | --- |
| AC_tissue_1 | -27.658456 | -56.074939 |  |
| AC_museum_1 | -27.612103 | -57.279819 | GPS location generalized Corrientes location |
| AC_museum_2 | -26.942803 | -54.516458 | GPS location generalized Misiones location |
| AC_museum_3 | -27.612103 | -57.279819 | GPS location generalized Corrientes location |
| AC_tissue_2 | -26.42362 | -53.83252 |  |
| AC_tissue_3 | -27.5275 | -55.867222 |  |
| AC_tissue_4 | -27.657928 | -56.075358 |  |
| AC_tissue_5 | -27.658442 | -56.075425 |  |
| AC_tissue_6 | -27.6584 | -56.0754 |  |
| AC_fecal_1 | -26.425012 | -53.835272 |  |
| AGC_museum_1 | -26.942803 | -54.516458 | GPS location generalized Misiones location |
| AGC_fecal_1 | -26.431315 | -53.836815 |  |
| AGC_fecal_2 | -26.431315 | -53.836815 |  |

### 4.3 Sample verification

We genotyped all samples collected from living *Alouatta* using published microsatellite protocols (Api06, Apm 01, and Apm04) (Cortés-Ortiz, Mondragón, & Cabotage, 2010) to ensure that each was collected from a unique individual. We separated PCR products on an 8% acrylamide gel (**Figure S1**) to determine microsatellite allele sizes.

To determine the sex of the museum and fecal samples, we amplified the SRY region using published protocols (Cortés-Ortiz et al., 2007; Di Fiore, 2006). We used Sanger sequencing (University of Utah Sequencing Core) on the SRY products. We considered samples to come from males if we successfully recovered SRY sequences with no ambiguities in the chromatograms.

Due to the age of the museum tissue samples, we measured DNA damage using mapDamage2.0 software (Jónsson, Ginolhac, Schubert, Johnson, & Orlando, 2013). DNA damage results in deamination of cytosine and guanine resulting in an excess of tyrosine and adenine substitutions, especially at the ends of sequencing reads (Briggs et al., 2007).

### 4.4 Shotgun sequencing and data processing

We shotgun sequenced the tissue sample with the highest concentration of DNA (AC_tissue_l) to create a reference for TLR7 (exons 1, 2, and 3), TLR8 (exons 1 and 2) and mtDNA genes ND1, ND2, and CO1. We sequenced mtDNA regions in order to compare genetic diversity at other loci to TLR7 and TLR8. The University of Utah Huntsman Cancer Center Sequencing Core prepared and shotgun sequenced the samples on an Illumina HiSeq 2500. We used BWA software (Li, 2013) to align *Alouatta* reads to the human reference genome (hg19) (1000 Genomes Project Consortium et al., 2015) and an unpublished *A. palliata* reference genome (A. Burrell, personal communication, 2018). Aligned BAM files were filtered to include only reads of MAPQ > 60 (Li, 2011). We used SPAdes software to create a consensus sequence for *Alouatta* TLR7 and TLR8 from the alignment (Bankevich et al., 2012) (**Supplementary Methods**).

### 4.5 Targeted sequencing

Using consensus TLR7, TLR8, ND1, ND2, and CO1 sequences we created a custom Ampliseq library (IAD149391-168, LifeTechnologies, Austin, TX) to perform targeted sequencing on the remainder of our samples. The University of Utah Sequencing Core completed targeted sequencing of the *A. caraya* and *A. guariba clamitans* samples using an Ion PGM and standard library kit protocols. To process the sequencing reads, we first trimmed the ends of the targeted sequencing data to remove erroneous sequencing and filtered fastq reads to maq > 20 using BBDuk (http://jgi.doe.gov/data-and-tools/bb-tools/). Second, we aligned trimmed fastq reads for each sample to the *A. caraya* TLR7, TLR8, ND1, ND2, and CO1 reference sequences using BWA software (Li, 2013). We used Freebayes software (Garrison & Marth, 2012) to jointly call variants in each species separately, and then merged the VCF files for analysis with bcftools (Li et al., 2009). We used vcftools to remove indels and filter all sites with GQ < 30 (Danecek et al., 2011). We omitted samples with greater than 40% data missing for TLR genes. After removing samples with too much missing data we removed sites where all samples were monomorphic or all data was missing. Finally, we excluded variable sites where more than six of the thirteen final samples (**Table 2**) were missing data. GenBank (Clark, Karsch-Mizrachi, Lipman, Ostell, & Sayers, 2016) accession numbers for all sequences can be found in **Table S1**.

**Table 2:**
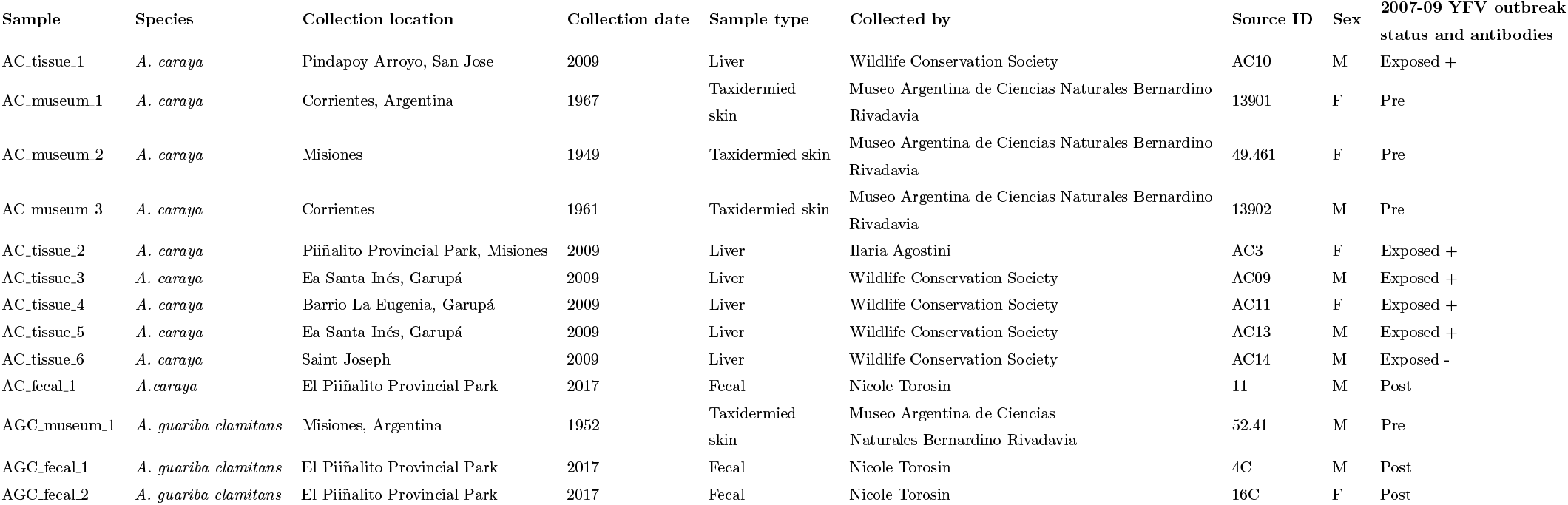
Samples

TLR7 and TLR8 are on the X chromosome in humans (Shen et al., 2010). However, *Alouatta* have multiple sex chromosomes. *A. caraya* females have four X chromosomes and males have two X and two Y chromosomes, while *A. guariba clamitans* females have six X chromosomes and males have three X chromosomes and two Y chromosomes (Steinberg, Nieves, & Mudry, 2014). Therefore, because males in both species have at least two X chromosomes, we did not correct male genotypes to be haploid, as is commonly done with human data. FreeBayes assumes that the genome is diploid when calling variants (Garrison & Marth, 2012). However, given that TLR7 and TLR8 are highly conserved (coding region identity between humans and *Alouatta*: TLR7 95.8%, TLR8 100%) we did not change the ploidy settings.

### 4.6 Analysis

We conducted principal component analyses (PCA) using FlashPCA (Abraham, Qiu, & Inouye, 2017) to assess how our samples clustered based on mtDNA and TLR data separately. We used DnaSP6 (Rozas et al., 2017) to measure nucleotide diversity (*π*) (Nei, 1987) within each species and Dxy, the mean number of pairwise differences (Takahata & Nei, 1985), between the following groups: *A. caraya* samples “Pre”, “Exposed”, and “Post” and *A. guariba clamitans* “Pre” and “Post” (**Table 3, Table 2**). DnaSP6 only considers sites without missing data in analyses (Rozas et al., 2017). We explored genetic differences between the exposure groups, revealed by PCA and Dxy, for functional implications.

**Table 3:** Summary of samples used in analyses.

| Species | Pre-outbreak (N) | Exposed to YFV and YFV antibody status | Alive post-outbreak |
| --- | --- | --- | --- |
| *A. caraya* | 3 | (+) = 5<br>(-) = 1 | 1 |
| *A. guariba clamitans* | 1 | 0 | 2 |

We compared the coding region of *A. guariba clamitans* and *A. caraya* with other primates of various YFV susceptibility. We obtained coding regions of TLR7 and TLR8 for New World and Old World primates from GenBank and dnazoo.org (Dudchenko et al., 2017) (**Table S2**). A. Burrell provided TLR7 and TLR8 sequences for *A. palliata*. We aligned DNA sequences using MEGA7 (S. Kumar, Stecher, & Tamura, 2016) and confirmed protein translation of TLR7 and TLR8 using reference NP_057646.1 and NP_619542.1, respectively. We aligned and edited TLR7 and TLR8 protein translations for all species using Aliview (Larsson, 2014).

We evaluated nonsynonymous changes in the coding region in *A.guariba clamitans* and *A. caraya* compared to the other primates in the alignment to identify variants potentially important to the survival of the individuals alive post-YFV outbreak.

## 5 Results

### 5.1 Samples and sequences

Sighting of *A. caraya* at El Piñalito Provincial Park in 2017 was the first in eight years. Sightings of *A. guariba clamitans* were the first in three years (I. Agostini, personal communication, 2019). We saw two *A. caraya*, one male and one female. We saw two groups of *A. guariba clamitans*, one with six individuals, including a juvenile, and one with three individuals. Microsatellite analysis revealed that we obtained fecal samples from one *A. caraya* and two *A. guariba clamitans*.

We generated sequences for nuclear genes TLR7 exons 1 (221 bp), 2 (196 bp), and 3 (5635), and TLR8 exons 1 (201 bp) and 2 (4217 bp). Additionally, we generated sequences for mtDNA genes ND1 (1075 bp), ND2 (1145 bp), and CO1 (1742 bp). We confirmed sex for the thirteen individuals (**Table 3, Table 2**). Most samples contained some missing data due to age of the tissue and skin samples and DNA fragmentation in fecal samples (Frantzen, Silk, Ferguson, Wayne, & Kohn, 1998). We did not detect post-mortem DNA damage in the museum samples (**Table S3**).

### 5.2 PCA

We created principal component plots for nuclear and mtDNA genetic loci using variant files filtered for completely monomorphic sites and sites with > 45% missing data. In our mtDNA and TLR principal component analyses, individuals clustered by species, not by YFV exposure status (**Figure S2, Figure S3**).

### 5.3 Diversity and divergence within each species

Nucleotide diversity within each species, *A. caraya* and *A. guariba clamitans*, is low (**Table 4**). We observed no polymorphic sites in TLR7 and TLR8 within each species and very little variation at the mtDNA loci.

**Table 4:** Nucleotide diversity within *Alouatta* species at each gene.

| Species | Gene | Segregating sites<br>(w/o missing data) | $\pi$ |
| --- | --- | --- | --- |
| *A. caraya* | TLR7 | 0 | 0 |
|  | TLR8 | 0 | 0 |
|  | ND1 | 1 | 0.0003 |
|  | ND2 | 3 | 0.0009 |
|  | COI | 1 | 6.90E-05 |
| *A. guariba*<br>*clamitans* | TLR7 | 0 | 0 |
|  | TLR8 | 0 | 0 |
|  | ND1 | 2 | 0.0007 |
|  | ND2 | 2 | 0.0009 |
|  | COI | 0 | 0 |

Comparisons of TLR7 and TLR8 sequences between the exposure groups within each species resulted in a Dxy value of zero (or n/a when there was only one total segregating site between the two groups being analyzed) (**Table S4, Table S5**). Dxy values for mtDNA between the groups are low (**Table S4, Table S5**). Thus, our results do not support our hypothesis regarding genetic differences between exposure groups. There are no genetic differences in TLR7 and TLR8 in the YFV surviving *Alouatta* individuals compared to those alive prior to the YFV outbreak or those exposed to YFV.

### 5.4 Coding sequence variation between species

While we did not observe divergence among the YFV exposure groups within each species, TLR7 and TLR8 sequences differ between *A. guariba clamitans* and *A. caraya*. Moreover, a number of these differences result in nonsynonymous amino acid changes compared to other primates. We identified four nonsynonymous variant sites in *A. guariba clamitans* compared to other primates. Two of these sites cause a biochemical property difference in the amino acid sequence. Both of these sites have been found to be positively selected (Torosin et al., 2019) (**Table 5**). Four nucleotide variants in *A. guariba clamitans* result in a synonymous change (**Table S6**).

**Table 5:**
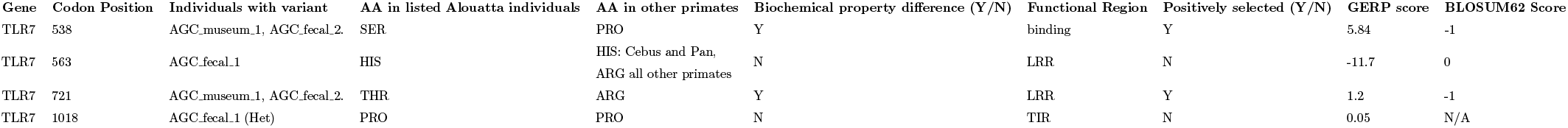
Variant sites, compared to other primates, in *A. guariba clamitans* samples with functional implications. Codon positions based on protein translation of TLR7, protein reference NP_057646.1.

All but one of the nucleotide differences (TLR7 codon 241) in *A. caraya* are synonymous (**Table S7**). The lone exception is a heterozygous variant in sample AC_tissue_2 at codon 241, resulting in an amino acid change.

We hypothesized that the surviving howler monkeys may possess advantageous genetic variants inherited from monkeys alive prior to the most recent YFV outbreak and that those that died of YFV in 2007-2009 may have lacked those variants. Alleles at greater frequency in *Alouatta* individuals alive after the 2007-2009 YFV outbreak may have been key in the individual survival for this extremely susceptible species.

## 6 Discussion

Our results did not support our hypothesis that surviving howler monkeys possess advantageous genetic variants at greater frequency than those alive before the YFV outbreak. Overall, we observed very little genetic variation within each *Alouatta* species. This is unsurprising for the highly conserved TLR7 and TLR8 and in line with results from other primates (Quach et al., 2013). The low diversity within the mtDNA is also expected. Limited polymorphism is thought to result from population expansion due to climatic changes at the start of the Holocene (Ascunce, Hasson, Mulligan, & Mudry, 2007). The mtDNA nucleotide diversity in our study is even lower than in previous studies despite the geographic and temporal variation of our samples. The small sample size and use of coding regions for analysis rather than the control region may be responsible for this.

Despite a lack of polymorphism within species, we identified nonsynonymous differences between species. Of the three nonsynonymous substitutions we identified in TLR7 of *A. guariba clamitans*, we found evidence that two are under positive selection (TLR7 codons 538 and 721) (Torosin et al., 2019) (**Table 5**). These two variants were identified in all *A. guariba clamitans* in this study and are unique to the species. The variants are within the extra-compartmental LRR region important in pathogen detection (H. Kumar, Kawai, & Akira, 2011). TLR7 codon 538 is in the binding region of the protein and directly interacts with ssRNA viruses such as YFV (Wei et al., 2009). The amino acid change to serine from proline at TLR7 codon 538 results in a biochemical change. The GERP score of this site is 5.84, indicating that it is extremely constrained throughout evolution (Cooper et al., 2005) and the BLOSUM62 score is −1, meaning that this substitution is not expected by chance (Henikoff & Henikoff, 1992). It is noteworthy that within a functionally important and evolutionarily constrained region, there is an amino acid change present in one species and the site is under positive selection solely in that species. The amino acid change at TLR7 codon 721 in *A. guariba clamitans* is also unique to the species. While the GERP score is lower, 1.2, it is still positive and has a BLOSUM score of −1, indicating evolutionary constraint at this locus (Henikoff & Henikoff, 1992). One of the other variants in *A. guariba clamitans* (TLR7 codon 563) results in an amino acid that is present in YFV resistant genera such as *Cebus*.

*A. caraya* shows only nonsynonymous changes within TLR7 and TLR8. One anomaly is AC_tissue_2. This is an interesting sample due to numerous heterozygous variants within the third exon of TLR7. This sample is from one of the tracked *Alouatta* groups (Agostini et al., 2010) in El Parque Piñalito that died from YFV (Holzmann et al., 2010). One of the heterozygous variants results in an amino acid that is fixed in all other New World primates (glycine). The second variant results in the amino acid that is fixed in Old World primates (aspartic acid). Neither pre-YFV museum samples nor YFV surviving samples possessed these variants. More samples from *A. caraya* from El Parque Piñalito are necessary to determine if these observations are unique to our study group.

It is noteworthy that we identified two nonsynonymous positively selected substitutions within a gene under evolutionary constraint that are unique to the *A. guariba clamitans* lineage. The genus *Alouatta* has the greatest range of all Neotropical primates (Crockett, 1998). *A. caraya* resides in dry semi-deciduous and deciduous forests in Brazil, Bolivia, Paraguay and Argentina (Crockett, 1998). *A. guariba clamitans* prefers a wetter, evergreen environment spanning the southern Atlantic coast of Brazil into Northern Argentina (Crockett, 1998). While sympatric in El Parque Piñalito today, prior to the 1940s, the two species were not sympatric, adhering to their preferred habitats (Aguiar et al., 2007). Potentially, the unique, positively selected genetic variants found in *A. guariba clamitans* may have resulted from specialized immune gene evolution for their particular environmental niche. This would be consistent with recent research showing that broadly conserved virus-interacting proteins, such as TLR7, are subject to strong adaptive events shaped by viruses (Castellano, Uricchio, Munch, & Enard, 2019). One study suggested that much of the current variability in *Alouatta* populations is due to recent events such as habitat alteration and disease (Chapman & Balcomb, 1998).

Extending analyses of TLR7 and TLR8 to a greater range of *Alouatta* species is essential to determine whether other species have the same amino acid substitutions identified here in *A. guariba clamitans*. More research is necessary to discover whether the genetic changes in *A. guariba clamitans* resulted from species-specific, environment specific pathogen-host interaction from the past, or whether they are from more recent changes in habitat and new diseases, such as YFV. There have been ongoing outbreaks of YFV in southeastern Brazil over the last two decades (Possas, Martins, Oliveira, & Homma, 2018) resulting in mass deaths of the species *A. guariba clamitans* (Centro de operações de emergências em saú pública sobre febre amarela, Ministerio de Saúde, 2017; Romano et al., 2014). It would be especially pertinent to obtain samples from this population to further investigate the expanse of these genetic variants in the *A. guariba clamitans* population and whether they may contribute to YFV resistance.

Results from this study warrant further investigation. The sample set available for this study was small and the biological material was often degraded resulting in missing data. More samples should be collected from the two *Alouatta* species in this study, as well as from *Alouatta* species across Central and South America, to fully explore the immune gene variation of the genus. In addition, expanding this work to include additional immune genes will be critical for fully understanding the potential genetic mechanisms underlying YFV susceptibility.

## 7 Acknowledgements

We acknowledge Patricia Casco and Candelaria Sanchez Fernandez for collection of the fecal samples. We would like to acknowledge the curators at the Museo Argentina de Ciencias Naturales Bernardino Rivadavia for the museum tissue samples. We would like to acknowledge Ines Badano and the members of her laboratory for their assistance in Argentina. We acknowledge the people of the Ministerio de Ecología y Recursos Naturales Renovables de Misiones Provincia for processing permits. We would like to thank Mike Powers and Derek Warner at the University of Utah Sequencing Core for assistance planning and implementing the seqeuncing protocols. We would like to thank Andy Burrell, Christina Bergey, and Todd Disotell for allowing us to use the *A. palliata* reference genome. Funding for this project was provided by the Center for Global Change and Sustainability at the University of Utah and the Wildlife Conservation Society.

## Supplementary Material

**Figure S1:**
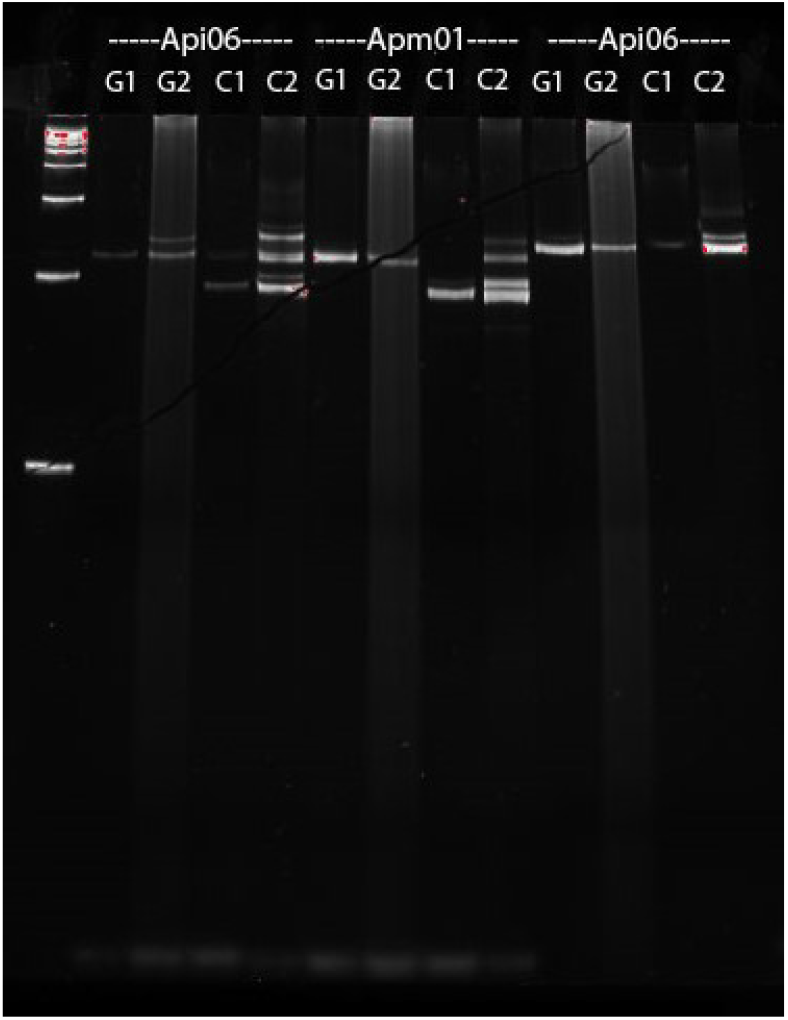
Microsatellite results to determine uniqueness of fecal samples collected from *A. guariba clamitans* (G) and *A. caraya* (C). Different alleles observed at Api06 for *A. guariba clamitans*. G1 (AGC_fecal_1) and G2 (AGC_fecal_2) are also male and female, respectively. Extra bands seen in the C2 lanes are due to non-specific amplification. *A. caraya samples* have the same alleles at all three loci.

**Figure S2:**
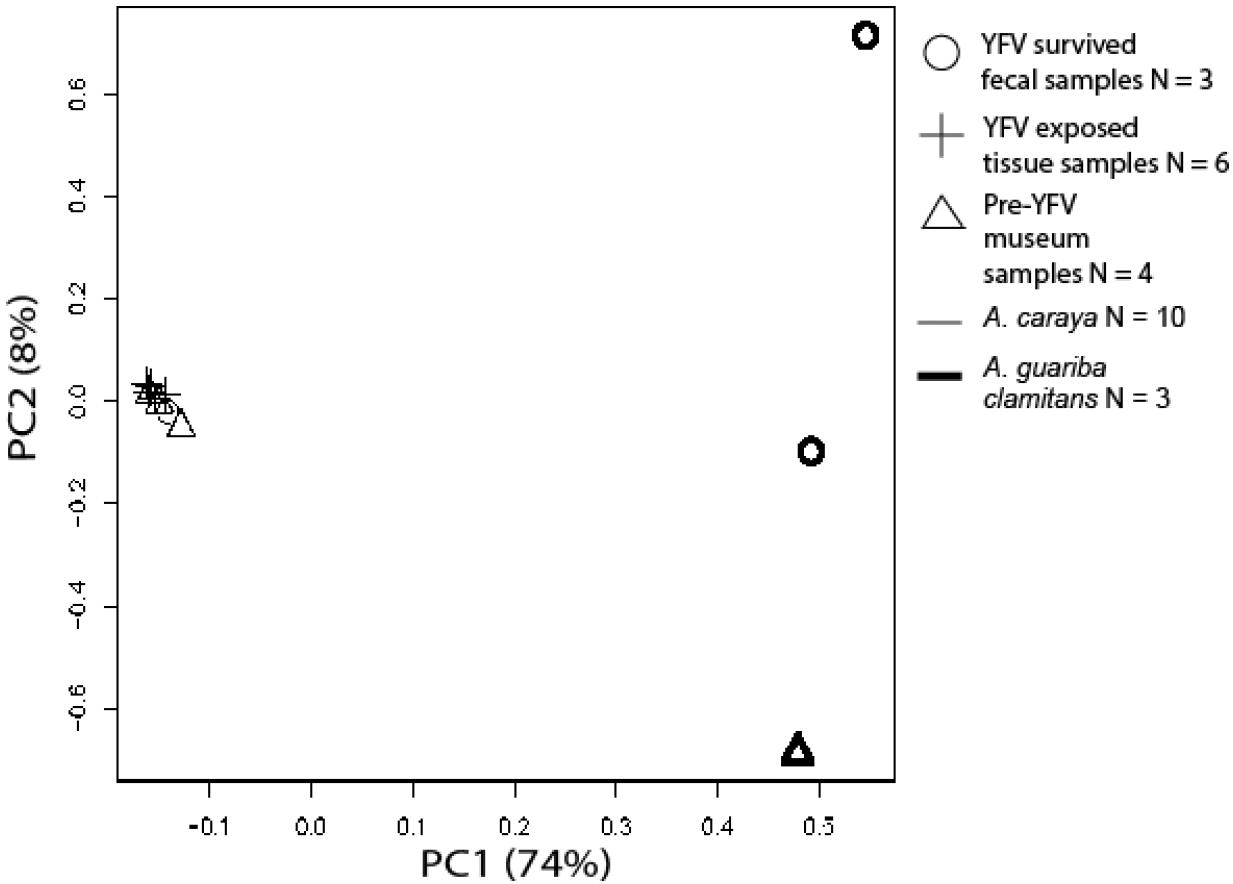
PCA analysis using variants from all *Alouatta* samples at all mtDNA genetic loci: ND1, ND2, CO1.

**Figure S3:**
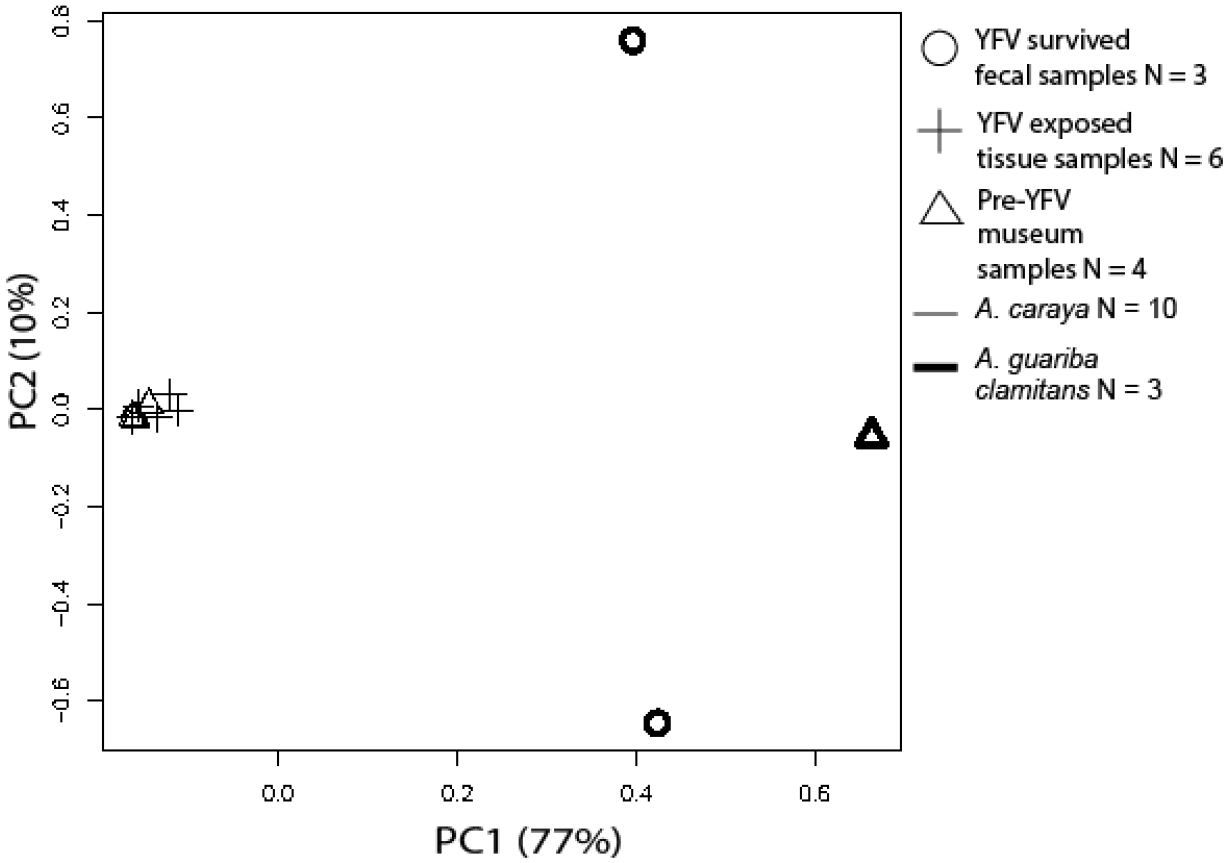
PCA analysis using variants from all *Alouatta* samples at all TLR7 and TLR8 genetic loci.

**Table S1:** GenBank Accession Numbers

| Sample | Gene | GenBank accession | Sample | Gene | GenBank accession |
| --- | --- | --- | --- | --- | --- |
| AGC_fecal_2 | TLR7 | MN235741 | AGC_fecal_1 | ND1 | MN235771 |
| AC_fecal_1 | TLR7 | MN235742 | AGC_fecal_2 | ND1 | MN235772 |
| AC_tissue_1 | TLR7 | MN046896 | AC_fecal_1 | ND1 | MN235773 |
| AC_tissue_2 | TLR7 | MN235743 | AC_tissue_1 | ND1 | MN046903 |
| AC_tissue_3 | TLR7 | MN235745 | AC_tissue_2 | ND1 | MN235774 |
| AC_tissue_4 | TLR7 | MN235746 | AC_tissue_3 | ND1 | MN235775 |
| AC_tissue_5 | TLR7 | MN235747 | AC_tissue_4 | ND1 | MN235776 |
| AC_tissue_6 | TLR7 | MN235748 | AC_tissue_5 | ND1 | MN235777 |
| AC_museum_1 | TLR7 | MN046898 | AC_tissue_6 | ND1 | MN235778 |
| AC_museum_2 | TLR7 | MN235749 | AC_museum_1 | ND1 | MN046906 |
| AC_museum_3 | TLR7 | MN235750 | AC_museum_2 | ND1 | MN235779 |
| AGC_museum_1 | TLR7 | MN072369 | AC_museum_3 | ND1 | MN235780 |
| AGC_fecal_1 | TLR8 | MN235751 | AGC_museum_1 | ND1 | MN046909 |
| AGC_fecal_2 | TLR8 | MN235752 | AGC_fecal_1 | CO1 | MN235781 |
| AC_fecal_1 | TLR8 | MN235753 | AGC_fecal_2 | CO1 | MN235782 |
| AC_tissue_1 | TLR8 | MN046897 | AC_fecal_1 | CO1 | MN235783 |
| AC_tissue_2 | TLR8 | MN235754 | AC_tissue_2 | CO1 | MN235784 |
| AC_tissue_3 | TLR8 | MN235755 | AC_tissue_3 | CO1 | MN235785 |
| AC_tissue_4 | TLR8 | MN235756 | AC_tissue_4 | CO1 | MN235786 |
| AC_tissue_5 | TLR8 | MN235757 | AC_tissue_5 | CO1 | MN235787 |
| AC_tissue_6 | TLR8 | MN235758 | AC_tissue_6 | CO1 | MN235788 |
| AC_museum_1 | TLR8 | MN046899 | AC_museum_1 | CO1 | MN046907 |
| AC_museum_2 | TLR8 | MN235759 | AC_museum_2 | CO1 | MN235789 |
| AC_museum_3 | TLR8 | MN235760 | AC_museum_3 | CO1 | MN235790 |
| AGC_museum_1 | TLR8 | MN046901 | AGC_museum_1 | CO1 | MN045881 |
| AGC_fecal_1 | ND2 | MN235761 |  |  |  |
| AGC_fecal_2 | ND2 | MN235762 |  |  |  |
| AC_fecal_1 | ND2 | MN235763 |  |  |  |
| AC_tissue_1 | ND2 | MN046902 |  |  |  |
| AC_tissue_2 | ND2 | MN235764 |  |  |  |
| AC_tissue_3 | ND2 | MN235765 |  |  |  |
| AC_tissue_4 | ND2 | MN235766 |  |  |  |
| AC_tissue_5 | ND2 | MN235767 |  |  |  |
| AC_tissue_6 | ND2 | MN235768 |  |  |  |
| AC_museum_1 | ND2 | MN046905 |  |  |  |
| AC_museum_2 | ND2 | MN235769 |  |  |  |
| AC_museum_3 | ND2 | MN235770 |  |  |  |
| AGC_museum_1 | ND2 | MN046908 |  |  |  |

**Table S2:**
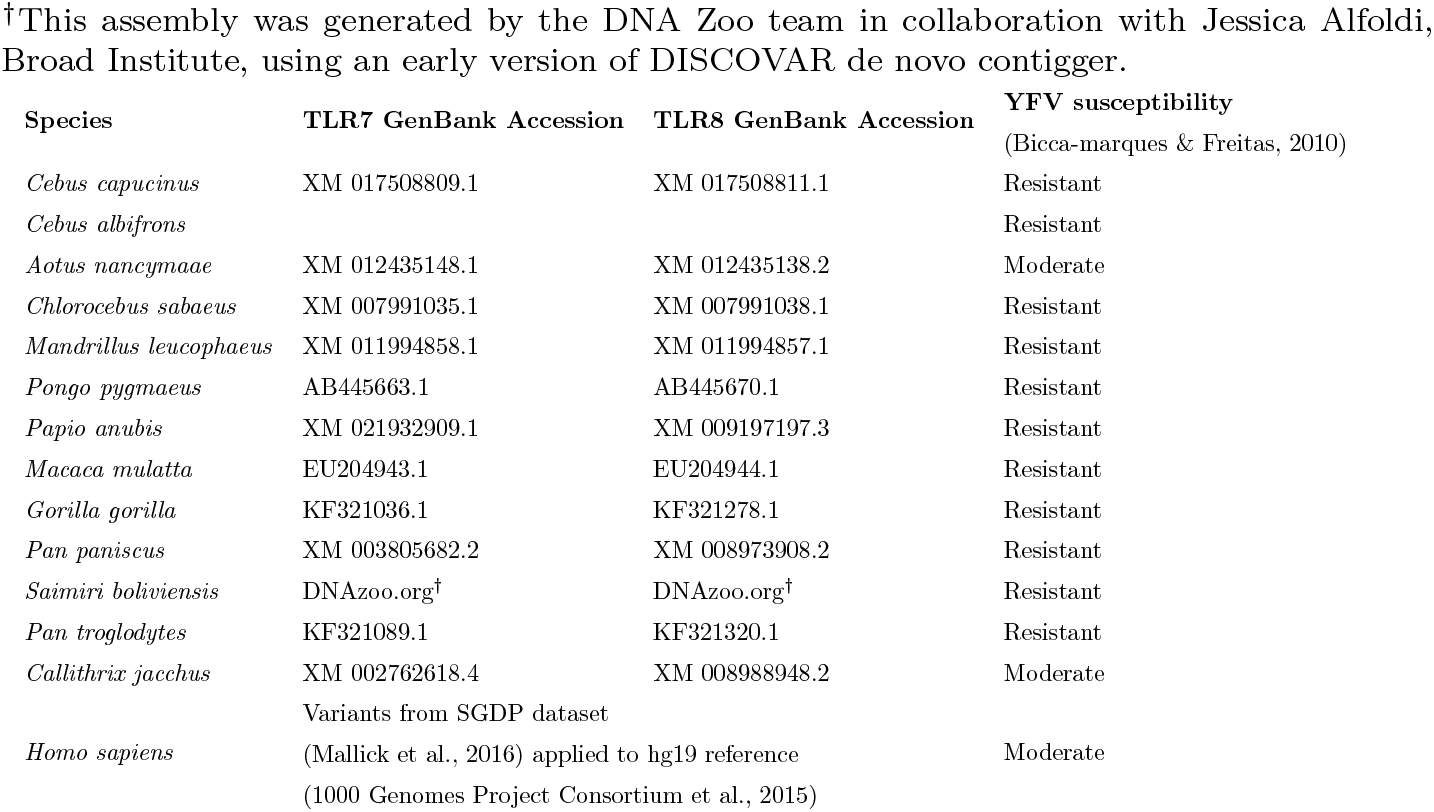
GenBank Accession numbers for TLR7 and TLR8 sequences and primate YFV susceptibility.

**Table S3:** MapDamage2.0 results for museum samples. Values indicate the difference in C>T substitutions between the sample and the reference at the first 5’ position. Values less than 0.01 indicate that level of DNA damage is low.

| Sample | Output |
| --- | --- |
| AC_museum_1 | 0.00374 |
| AC_museum_2 | 0.00480 |
| AC_museum_3 | 0.00328 |
| AGC_museum_1 | 0.00485 |

**Table S4:** DnaSP results for *A. caraya* variant data. Only sites without missing data included in analysis. Pop1: AC_museum_1, AC_museum_2, AC_museum_3 Pop2: AC_tissue_1, AC_tissue_2, AC_tissue_3, AC_tissue_4, AC_tissue_5, AC_tissue_6 Pop3: AC_fecal_1

| Population_1 | Pop_2 | Loci | Total Differences | Dxy |
| --- | --- | --- | --- | --- |
| 2 | 1 | TLR7e2 | 0 | n/a |
| 2 | 3 | TLR7e2 | 0 | n/a |
| 1 | 3 | TLR7e2 | 0 | n/a |
| 2 | 1 | TLR7e3 | 0 | 0 |
| 2 | 3 | TLR7e3 | 0 | 0 |
| 1 | 3 | TLR7e3 | 0 | 0 |
| 2 | 1 | TLR8e2 | 0 | 0 |
| 2 | 3 | TLR8e2 | 0 | 0 |
| 1 | 3 | TLR8e2 | 0 | 0 |
| 2 | 1 | TLR8e1 | 0 | n/a |
| 2 | 3 | TLR8e1 | 0 | n/a |
| 1 | 3 | TLR8e1 | 0 | n/a |
| 2 | 1 | ND2 | 5 | 0.0022 |
| 2 | 3 | ND2 | 3 | 0.0021 |
| 1 | 3 | ND2 | 3 | 0.0015 |
| 2 | 1 | ND1 | 2 | 0.00075 |
| 2 | 3 | ND1 | 2 | 0.00063 |
| 1 | 3 | ND1 | 1 | 0.00036 |
| 2 | 1 | CO1 | 0 | 0 |
| 2 | 3 | CO1 | 1 | 0.00035 |
| 1 | 3 | CO1 | 2 | 0.000578 |

**Table S5:** DnaSP results for *A. guariba clamitans* variant data. Only sites without missing data included in analysis. Pop1: AC_museum_1, AC_museum_2, AC_museum_3 Pop2: AC_tissue_1, AC_tissue_2, AC_tissue_3, AC_tissue_4, AC_tissue_5, AC_tissue_6 Pop3: AC_fecal_1

| Population_1 | Pop_2 | Loci | Total Differences | Dxy |
| --- | --- | --- | --- | --- |
| 1 | 2 | TLR7e2 | 0 | n/a |
| 1 | 2 | TLR7e3 | 0 | 0 |
| 1 | 2 | TLR8e2 | 0 | 0 |
| 1 | 2 | TLR8e1 | 0 | n/a |
| 1 | 2 | ND2 | 0 | 0.00082 |
| 1 | 2 | ND1 | 2 | 0.0012 |
| 1 | 2 | CO1 | 0 | 0 |

**Table S6:**
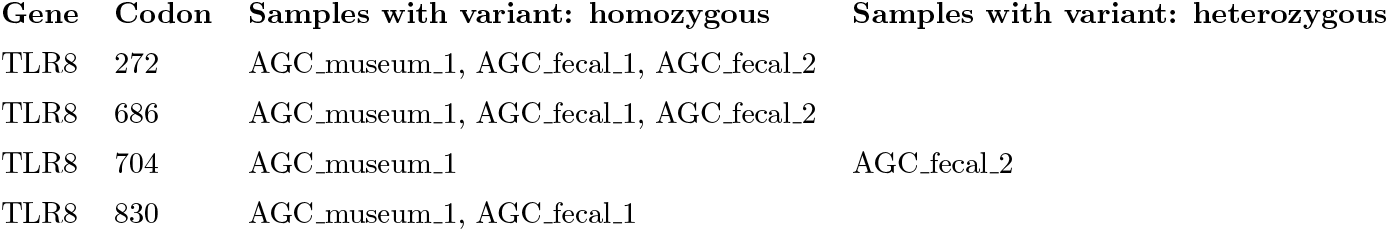
Synonymous variant loci in *A. guariba clamitans*.

**Table S7:**
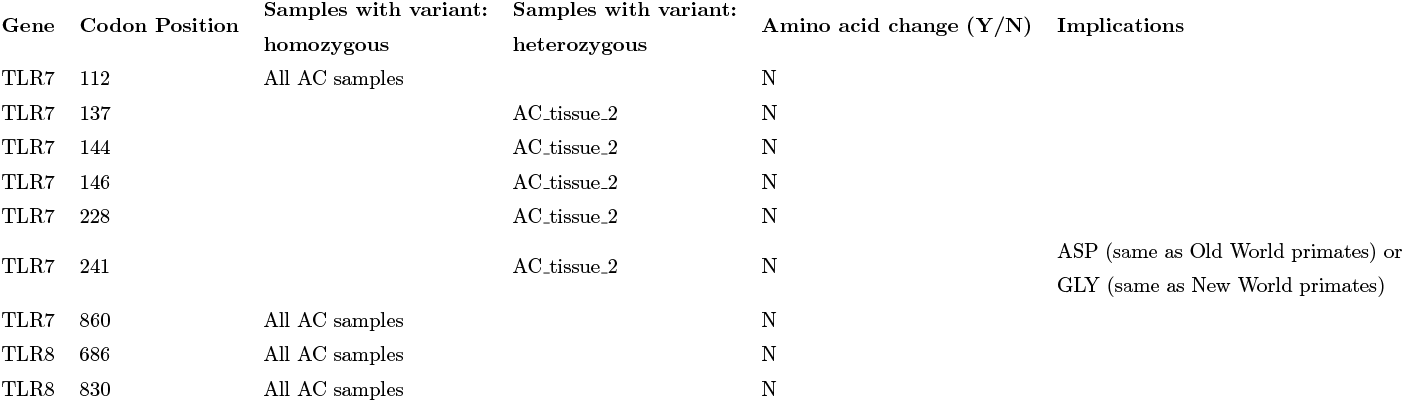
*A. caraya* variants: Variant sites in *A. caraya* samples with functional implications. Codon positions based on protein translation of TLR8, protein reference NP619542.1.

| Gene | Codon Position | Samples with variant: homozygous | Samples with variant: heterozygous | Amino acid change (Y/N) | Implications |
| --- | --- | --- | --- | --- | --- |
| TLR7 | 112 | All AC samples |  | N |  |
| TLR7 | 137 |  | AC.tissue_2 | N |  |
| TLR7 | 144 |  | AC.tissue_2 | N |  |
| TLR7 | 146 |  | AC.tissue_2 | N |  |
| TLR7 | 228 |  | AC.tissue_2 | N |  |
| TLR7 | 241 |  | AC.tissue_2 | N | ASP (same as Old World primates) or<br>GLY (same as New World primates) |
| TLR7 | 860 | All AC samples |  | N |  |
| TLR8 | 686 | All AC samples |  | N |  |
| TLR8 | 830 | All AC samples |  | N |  |

## 8 Methods

We processed targeted sequencing results using the following pipeline:

1. Unalign BAM files for AC_museum and AGC_museum using GATK software (Danecek et al., 2011).

~~~
java – jar /GATK_Resource/picard–tools–1.134/picard.jar RevertSam I={input.bam} O={output.unaligned_bam}
~~~
2. Convert unaligned BAM files to Fastq format (Quinlan, 2014).

~~~
bedtools bamtofastq −i {input.unaligned_bam} −fq {output.fastq}
~~~
3. Assess fastq quality. Software available at http://www.bioinformatics.babraham.ac.uk/projects/fastqc/

~~~
fastqc −o fastqc_results sample.fastq
~~~
4. Trim fastq reads based on fastq quality results. Software available at https://sourceforge.net/projects/bbmap/

~~~
bbduk.sh −Xmx1g in={input.fq} out={output.out_fq} ftl=10 ftr=200 minlen=75 maq=20
~~~
5. Realign trimmed Fastq files for each sample to AC_tissue reference (Li & Durbin, 2009; Li et al., 2009).

~~~
bwa mem −R @RG\\tID:{id}\\tSM:{id}\\tLB:{id}\\tPU:{id}\\ tPL:
{IonPGM} {input.ref} {input.fq} | samtools fixmate −O bam – – | samtools sort −O bam −o {output.bams}
~~~
6. Index each BAM file (Li et al., 2009).

~~~
samtools index {input}
~~~
7. Mark duplicate reads using GATK software (Danecek et al., 2011).

~~~
java −jar /GATK\_Resource/picard–tools–1.134/picard.jar MarkDuplicates I={input.bam} O={output.bam} M={output.metrics}
~~~
8. Create a consensus VCF based on the BAM files (with duplicate reads marked) of each species (Garrison & Marth, 2012).

~~~
freebayes ––fasta–reference AC_tissue.fasta ––report −monomorphic ––genotype–qualities ––L {list_of_bam_files} −v {output.vcf}
~~~
9. Remove samples with > 40% missing data within TLR genes.
10. Merge VCF files (Li et al., 2009).

~~~
bcftools merge {A_caraya.vcf.gz} {A_guariba_clamitans.vcf.gz} −O v −o {AC_AGC_merged.vcf}
~~~
11. Remove indels from VCF file and filter by genotype quality (Danecek et al., 2011). Indels were removed since they appeared to be related to sequencing error caused by long strings of nucleotide adenine.

~~~
vcftools ––vcf {AC_AGC_merged.vcf} ––minGQ 30 ––remove–indels ––recode −out \newline {AC_AGC_merged_minGQ30.vcf}
~~~
12. Remove variant sites where every sample is monomorphic or missing data.
13. Remove variant sites where >6 samples are missing data (out of 13 total).

